## Supplementary File 1 for "PI3K/HSCB axis facilitates FOG1 nuclear translocation to promote erythropoiesis and megakaryopoiesis"

**Supplementary File 1—table 1. Primary antibodies used in this study**

| **Antibody name** | **Catalog No. and vendor** | **Amount (dilution fold)** |
| --- | --- | --- |
| CD34 Monoclonal Antibody (4H11), FITC, eBioscience™ | 11-0349-42, Thermo Scientific™, USA | 0.50 µg/100 µL |
| CD90 Monoclonal Antibody (eBio5E10 (5E10)), APC, eBioscience™ | 17-0909-42, Thermo Scientific™, USA | 0.25 µg/100 µL |
| [CD71 Monoclonal Antibody (OKT9 (OKT-9)), APC, eBioscience™](https://www.thermofisher.cn/antibody/product/CD71-Transferrin-Receptor-Antibody-clone-OKT9-OKT-9-Monoclonal/17-0719-42) | 17-0719-42, Thermo Scientific™, USA | 0.10 µg/100 µL |
| CD235a Monoclonal Antibody (10F7MN), FITC, eBioscience™ | 11-9886-42, Thermo Scientific™, USA | 0.25 µg/100 µL |
| CD38 Monoclonal Antibody (HIT2), APC, eBioscience™ | 17-0389-42, Thermo Scientific™, USA | 0.25 µg/100 µL |
| CD45RA Monoclonal Antibody (HI100), FITC, eBioscience™ | 11-0458-42, Thermo Scientific™, USA | 1.50 µg/100 µL |
| CD123 (IL3RA) Monoclonal Antibody (6H6), APC, eBioscience™ | 17-1239-42, Thermo Scientific™, USA | 0.25 µg/100 µL |
| Monoclonal Mouse anti‑Human EPOR Antibody (clone 12K90, APC) | LS‑C182845, LifeSpan Biosciences, USA | 5 µL/100 µL |
| CD71 Monoclonal Antibody (OKT9 (OKT-9)), FITC, eBioscience™ | 11-0719-42, Thermo Scientific™, USA | 0.25 µg/100 µL |
| CD41a Monoclonal Antibody (HIP8), APC, eBioscience™ | 17-0419-42, Thermo Scientific™, USA | 0.25 µg/100 µL |
| CD42b Monoclonal Antibody (HIP1), FITC, eBioscience™ | 11-0429-42, Thermo Scientific™, USA | 1.00 µg/100 µL |
| Mouse anti DDDDK-Tag mAb | AE005, ABclonal, China | 1:100 (for IP) |
| Anti-TACC3 Antibody | ab245455, Abcam, UK | 1:50 (for IP) |
| FOG1/ZFPM1 Polyclonal Antibody | A301-431A-T, Thermo Scientific™, USA | 1:50 (for IP) |
| Anti-HSCB antibody produced in rabbit | HPA018447, Sigma-Aldrich, USA | 1:50 (for IP) |
| PI3K p85 alpha Monoclonal Antibody (U5) | MA1-74183, Thermo Scientific™, USA | 1:50 (for IP) |
| HSCB Mouse Monoclonal Antibody [Clone ID: OTI3E1] | TA507274, OriGene, USA | 1:1000 (for WB) |
| Anti-ISCU Antibody [OTI4F5] | ab180532, Abcam, UK | 1:2000 (for WB) |
| NFS1 Rabbit pAb | A13385, ABclonal, China | 1:1000 (for WB) |
| GAPDH Rabbit Polyclonal Antibody | LF206, Epizyme, China | 1:2000 (for WB) |
| Recombinant Anti-Glycophorin A Antibody [EPR8200] | ab129024, Abcam, UK | 1:1000 (for WB) |
| alpha 1 Spectrin Rabbit mAb | A9597, ABclonal, China | 1:1000 (for WB) |
| ZFPM1 Polyclonal Antibody | PA5-40500, Thermo Scientific™, USA | 1:1000 (for WB) |
| NAA30 Polyclonal Antibody | PA5-107003, Thermo Scientific™, USA | 1:1000 (for WB) |
| [KO Validated] LYAR Rabbit pAb | A19900, ABclonal, China | 1:1000 (for WB) |
| GATA1 Rabbit mAb | A21262, ABclonal, China | 1:1000 (for WB) |
| [KO Validated] Lamin B1 Rabbit pAb | A16909, ABclonal, China | 1:1000 (for WB) |
| β-actin Mouse Monoclonal Antibody | LF201, Epizyme, China | 1:2000 (for WB) |
| TACC3 Rabbit mAb | A19617, ABclonal, China | 1:2000 (for WB) |
| MYO1E Polyclonal Antibody | PA5-100624, Thermo Scientific™, USA | 1:1000 (for WB) |
| Phosphoserine/threonine/tyrosine Polyclonal Antibody | 61-8300, Thermo Scientific™, USA | 1:200 (for WB) |
| PI3 Kinase p85 alpha Rabbit mAb | A4992, ABclonal, China | 1:1000 (for WB) |
| STAT5 Rabbit mAb | A5029, ABclonal, China | 1:1000 (for WB) |
| Phospho-Stat5 (Tyr694) (D47E7) XP® Rabbit mAb | 4322, Cell Signalling Technology, USA | 1:1000 (for WB) |
| Akt (pan) (C67E7) Rabbit mAb | 4691, Cell Signalling Technology, USA | 1:1000 (for WB) |
| Phospho-Akt (Ser473) Antibody | 9271, Cell Signalling Technology, USA | 1:1000 (for WB) |
| MEK1/2 (L38C12) Mouse mAb | 4694, Cell Signalling Technology, USA | 1:1000 (for WB) |
| Phospho-MEK1/2 (Ser221) (166F8) Rabbit mAb | 2338, Cell Signalling Technology, USA | 1:1000 (for WB) |

**Supplementary File 1—table 2. Primers used in the qRT-PCR assays**

| **Primer** | **Sequence (5’→3’)** |
| --- | --- |
| Human *GAPDH* qF | TCATCAGCAATGCCTCCT |
| Human *GAPDH* qR | CATCACGCCACAGTTTCC |
| Human *HBB* qF | ATGAAGTTGGTGGTGAGG |
| Human *HBB* qR | TGAGGTTGTCCAGGTGAG |
| Human *ALAS2* qF | GAAGTCTAACCCTAAGATACCC |
| Human *ALAS2* qR | GACCCATACAGTCCTACAGC |
| Human *GYPC* qF | TGTGATTGCTGCTGTGGC |
| Human *GYPC* qR | TCTTGGAGGGCAGGGTCT |
| Human *VWF* qF | GGTGATCGCCTCTTATGC |
| Human *VWF* qR | CCTTCGGAGGGATGGTC |
| Human *GP1BA* qF | TCTACCTGAAAGGCAATGAG |
| Human *GP1BA* qR | TTGGAGGAGAAGGGTGTC |
